# Order of events in a developing genetic code

**DOI:** 10.1101/2022.12.31.522385

**Authors:** Michael Yarus

## Abstract

Preexisting partial genetic codes can fuse to evolve toward the Standard Genetic Code (SGC). Code fusion provides a path of least selection, generating a code precursor that resembles the SGC, consequently evolving quickly. Optimal evolution requires wobble coding delayed until late in primordial codon assignment, because early wobble specifically retards evolution of complete and accurate codes. Given delayed wobble, the SGC can emerge after a modest selection for more proficient encoding.

## Introduction

Recent analysis clarifies an evolutionary path to the Standard Genetic Code (SGC); it invokes only typical, explicit evolutionary events, yet quantitatively realizes the SGC within a feasible protobiotic context.

The proposed path (Yarus 2021b, 2022a) involves partial codes, formed without wobble, which are fused, combining independently arising coding compartments. Their coding may have solved different problems in their separate origins, but partial codes with compatible assignments can be combined to approach ‘complete’ coding (20 amino acids plus start and stop) and ‘full’ coding (all triplets assigned), as in the SGC.

Fusion of partially-assigned codes is pivotal. It gathers codes that function across a scattered, possibly divergent, population, even before a unified genome exists (Aggarwal et al. 2016; Froese et al. 2018). Moreover, it has notable corrective powers. A fusing population excludes deviation: for example, it excludes random assignments, or assignments unrelated to an underlying organizing principle (Yarus 2022a). Because only fused codes with compatible assignments will be highly fit, the population of successfully fused codes is more homogeneous than its separated precursors. A fusing population therefore converges to a unified, common coding scheme. If nucleated by stereochemical assignments consistent with the SGC, such a population evolves toward the SGC.

Therefore, an origin in which some cognate coding triplets existed in RNA binding sites for amino acids is plausible (Yarus 2017); these RNA sequences could serve as primordial coding triplets. It seems unlikely that all such RNA-amino acid interactions have been detected by modern RNA-amino acid affinity selections conducted under laboratory conditions. So, the initial RNA foundation for the SGC may be broader than currently apparent. However, because amino acids do not equally readily form RNA complexes, it is also plausible that some SGC assignments were made later, by other means; for example, when early amino acids or peptides collaborated with primordial RNA in coding complexes. As examples, such collaboration could originate with aminoacylated RNAs (Szathmáry 1993), peptidyl-RNAs formed on an oligonucleotide aminoacylation catalyst (Turk et al. 2011), noncovalent RNA-peptide complexes (Carter 2015) or perhaps primordial ribonucleoprotein granules (Ripin and Parker 2022). Continuing code fusion would build later ribonucleopeptide assignments on a pre-existing amino acid-RNA binding foundation, likely conserving earliest RNA assignments in the final, near-universal SGC.

## Results

### Varied wobble onset

Because this work varies onset times for simple wobble (Crick 1966), Fig. 1 portrays fraction of wobbling assignments when Pwob = 0.0, 0.0004, 0.001, 0.005 and 0.02/passage. A passage is the unit of evolutionary time – the interval for one step of code evolution (see Methods; Yarus 2021a). For example, Pwob = 0.001/passage yields a mean of 1 wobble initiation at 1000 passages. This implies that [1-e^-1^] = 63.2% of Pwob = 0.001 environments have begun wobble coding after 1000 passages, as for the blue curve in Fig. 1. Below, 63.2 % wobble occurs late (at infinity) down to earliest at 50 passages. Note especially that even with very slow wobble onset (e.g., 0.0004/passage in Fig. 1, red), significant commitment to wobble occurs early – realistic wobble appearance is not sudden, but obeys an exponential progression with some environments wobbling, even at early times.

**Figure 1.**
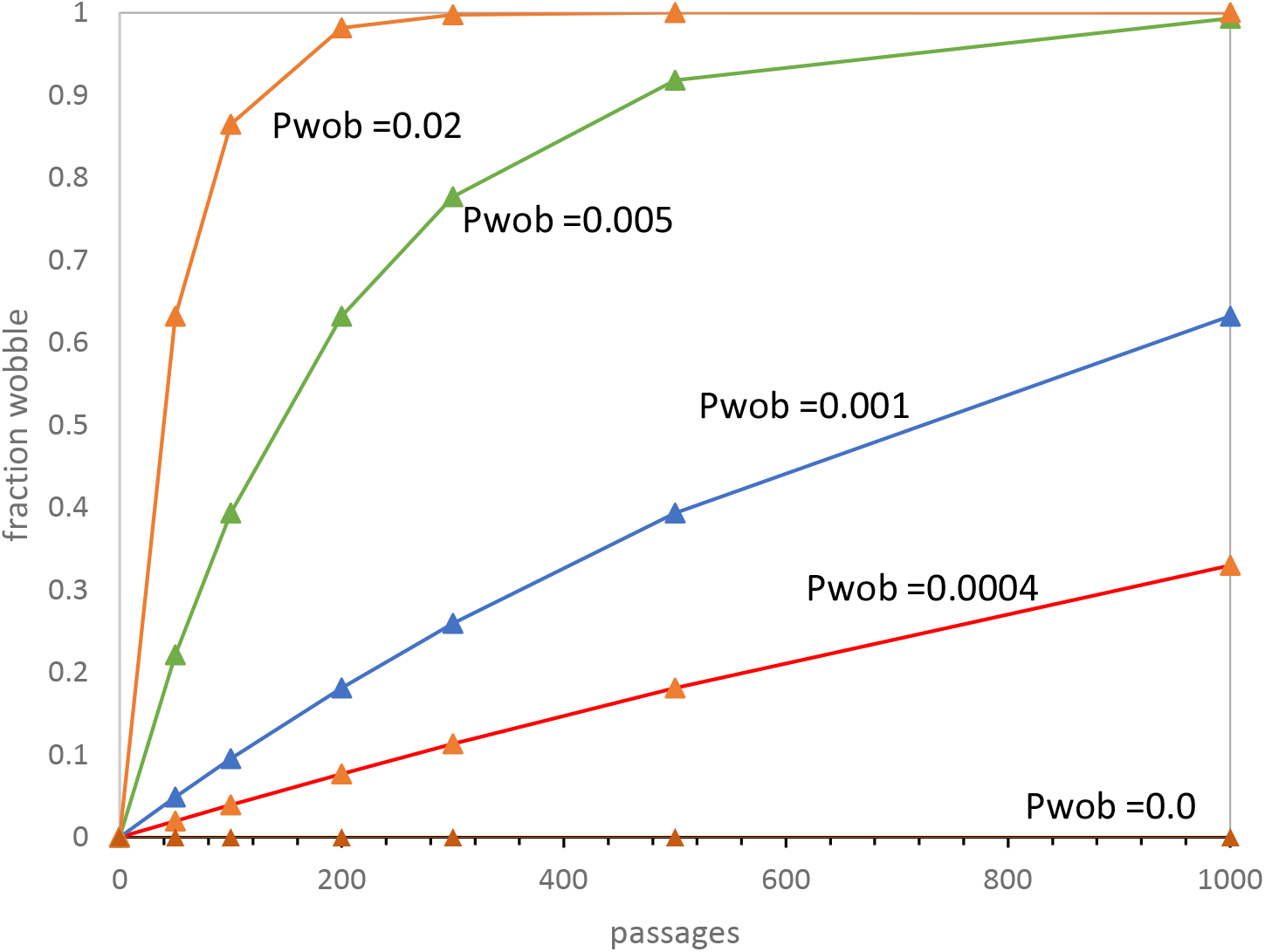
Progressive advent of wobble coding. Mean fraction of environmental codon assignments utilizing simple Crick wobble instead of standard base pairing is plotted versus time in passages. Plots cover times discussed in the text, for Pwob = 0.0 (brown), 0.0004 (red), 0.001 (blue), 0.005 (green) and 0.02 (orange) per passage. Other probabilities have standard values: Pmut (related assignment to unused neighbor) = 0.00975, Pdecay (loss of assignment) = 0.00975, Pinit (initial assignment) = 0.15, Prand (random assignment) = 0.05, Pfus (code fusion) = 0.002, Ptab (new code arises) = 0.08 per passage.

### Early wobble and complete coding

Early wobble is obstructive to code evolution (Fig. 2). With no wobble (Pwob = 0.0, wobble onset at indefinitely long time), mean time to evolve >= 20 encoded functions is about 670 passages. But if wobble begins earlier (Pwob = 0.01, 63.2% at 100 passages), near-complete coding is very significantly delayed, to about 3500 passages. Wobble hindrance increases approximately linearly for slowly-appearing wobble, but reaches a limit when wobble is essentially always present (rightward in Fig. 2). Thus, to investigate wobble’s negative effects, conditions at the left of Fig. 2 (no wobble, Pwob = 0.0) are compared with those rightward in Fig. 2 (wobble nearly always present, Pwob = 0.02).

**Figure 2.**
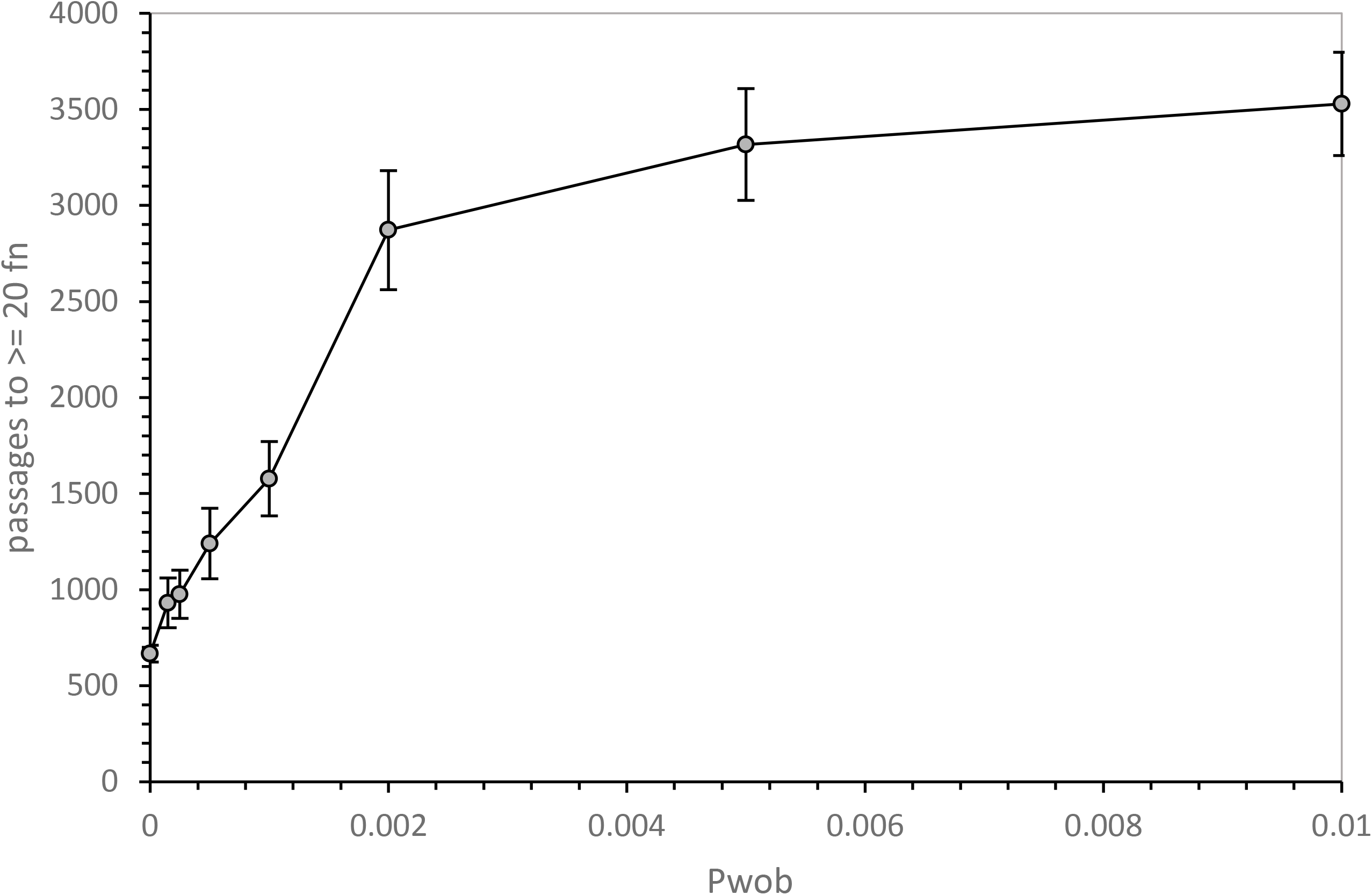
Early wobble delays complete coding. Pwob (as in Fig. 1) is plotted versus mean time (in Passages, for 100 environments) to evolve the first code with >= 20 assigned functions in an environment. System constants are those listed in Fig. 1. Error bars are standard errors of means.

### A time of optimal completeness

Fig. 3 begins to resolve time effects. It shows the fraction of active codes (that is, unfused + successfully fused codes) in 1000 evolving code environments, with (blue, Pwob = 0.02, c.f. Fig. 1) and without (red, Pwob = 0.0, Fig. 1) simple Crick wobble (Yarus 2021a). Under these conditions, a peak of near-complete coding systems (>= 20 fn encoded) transiently evolves near 300 passages (Fig. 1). But even more striking, early wobble suppresses complete coding (reduces it to ≈ 18%) without greatly changing its timing. Thus, Fig. 3 suggests an explanation for evolutionary delays in Fig. 2: wobble reduces the number of near-complete codes. Because near-complete coding systems are the probable precursors of the SGC, wobble hinders SGC evolution. Wobble inhibition of codes specifying >= 20 encoded functions takes the form of a reduced peak at similar times; wobble inhibition is not a time delay which will be corrected later.

**Figure 3.**
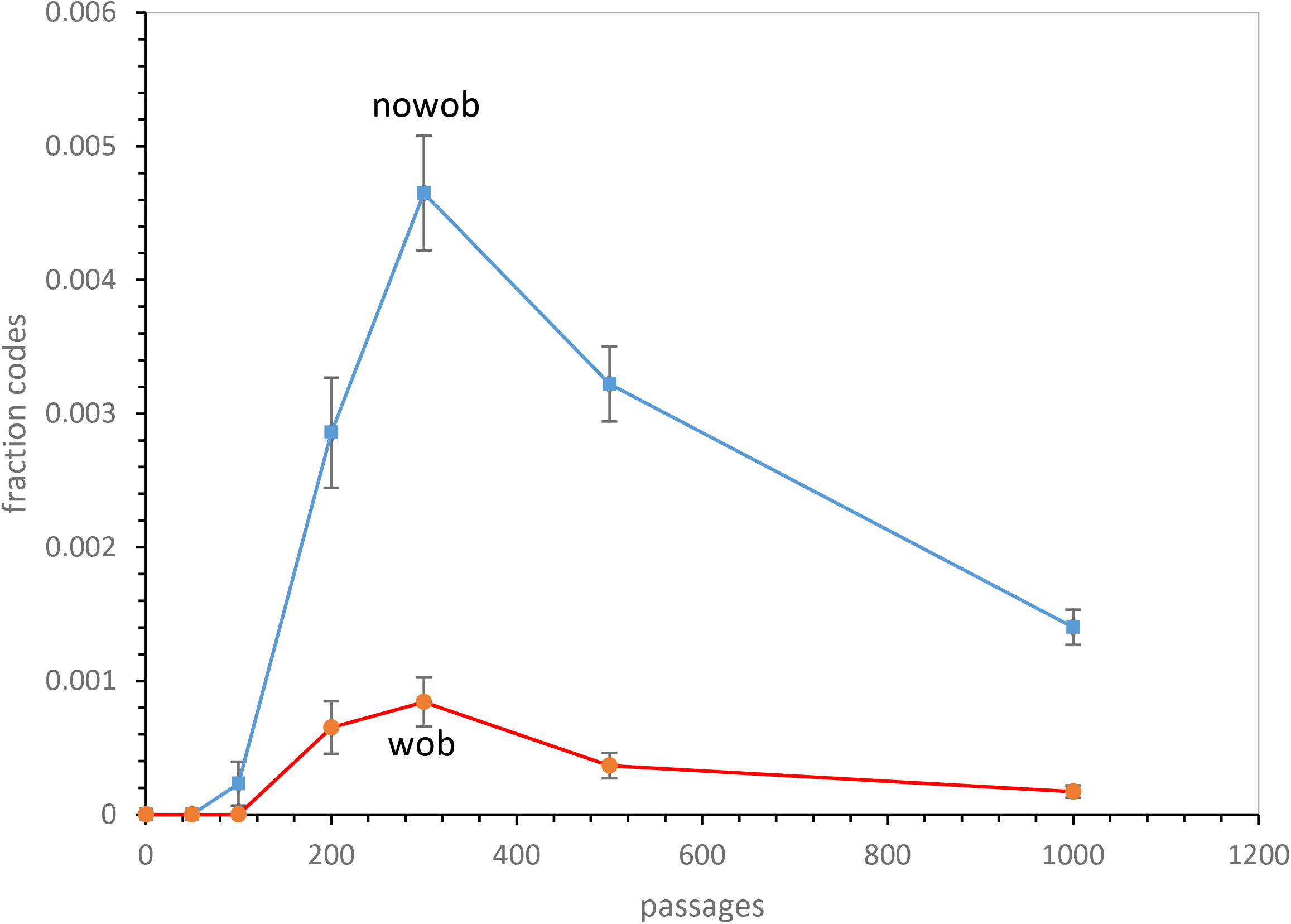
Early wobble obstructs complete coding. Fraction of active codes in 1000 environments that have >= 20 assigned functions vs time in passages, for Pwob = 0.0 (nowob, squares) and Pwob = 0.02 (wob, circles). except for Pwob, probabilities/passage are those in Fig. 1. Error bars are standard errors.

### Fused and unfused codes make distinguishable contributions

Fig. 4A shows the distribution of completion among codes active in peak environments (300 passages, Fig. 3). Fig. 4A plots the frequency of each such code versus its number of encoded functions. Active codes with only 2 functions are most frequent (blue, at left), and the relative abundance of more complete codes declines regularly, though codes with 20 and 21 functions exist at low levels among environments at 300 passages (blue, at right).

**Figure 4.**
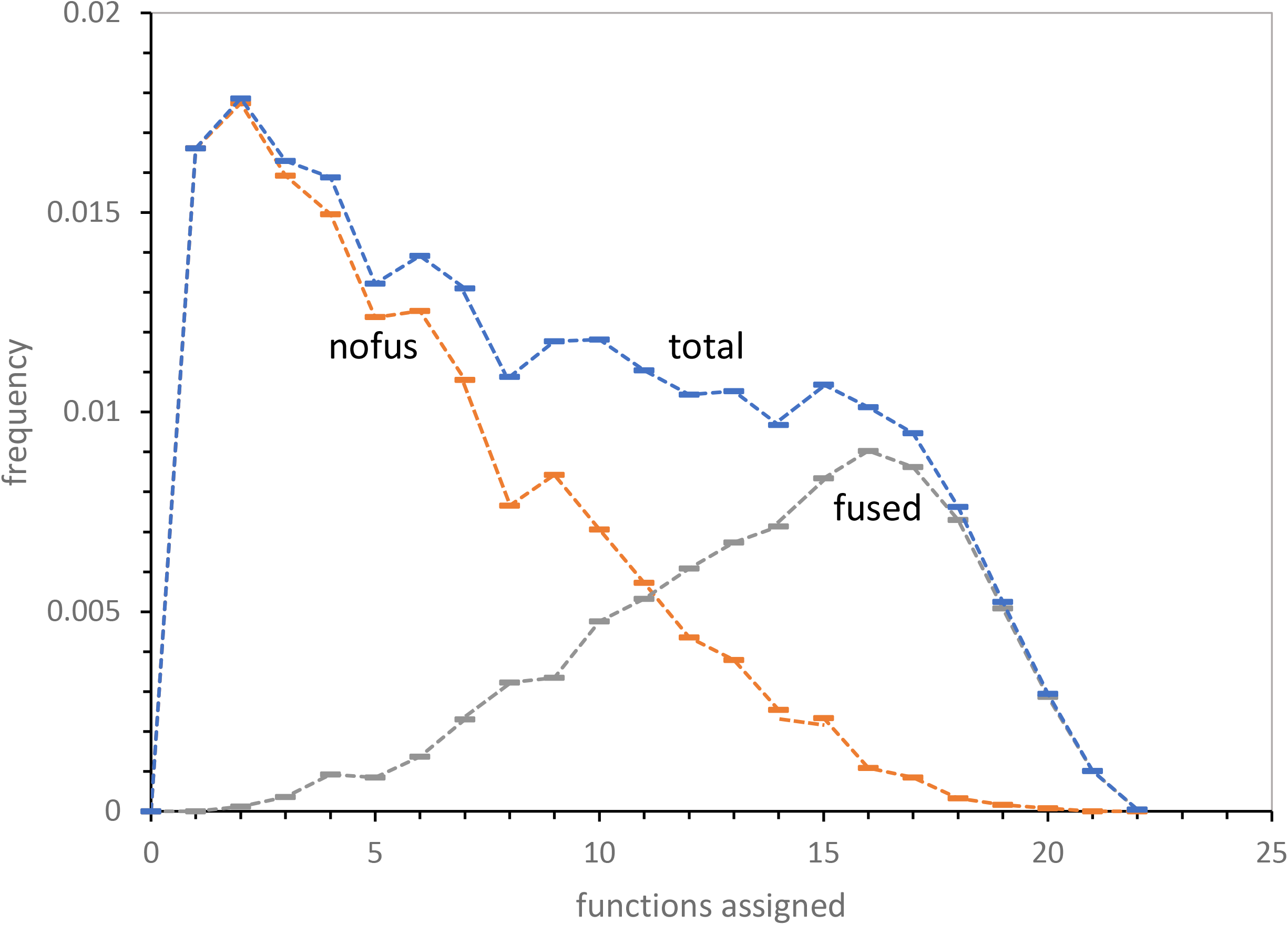
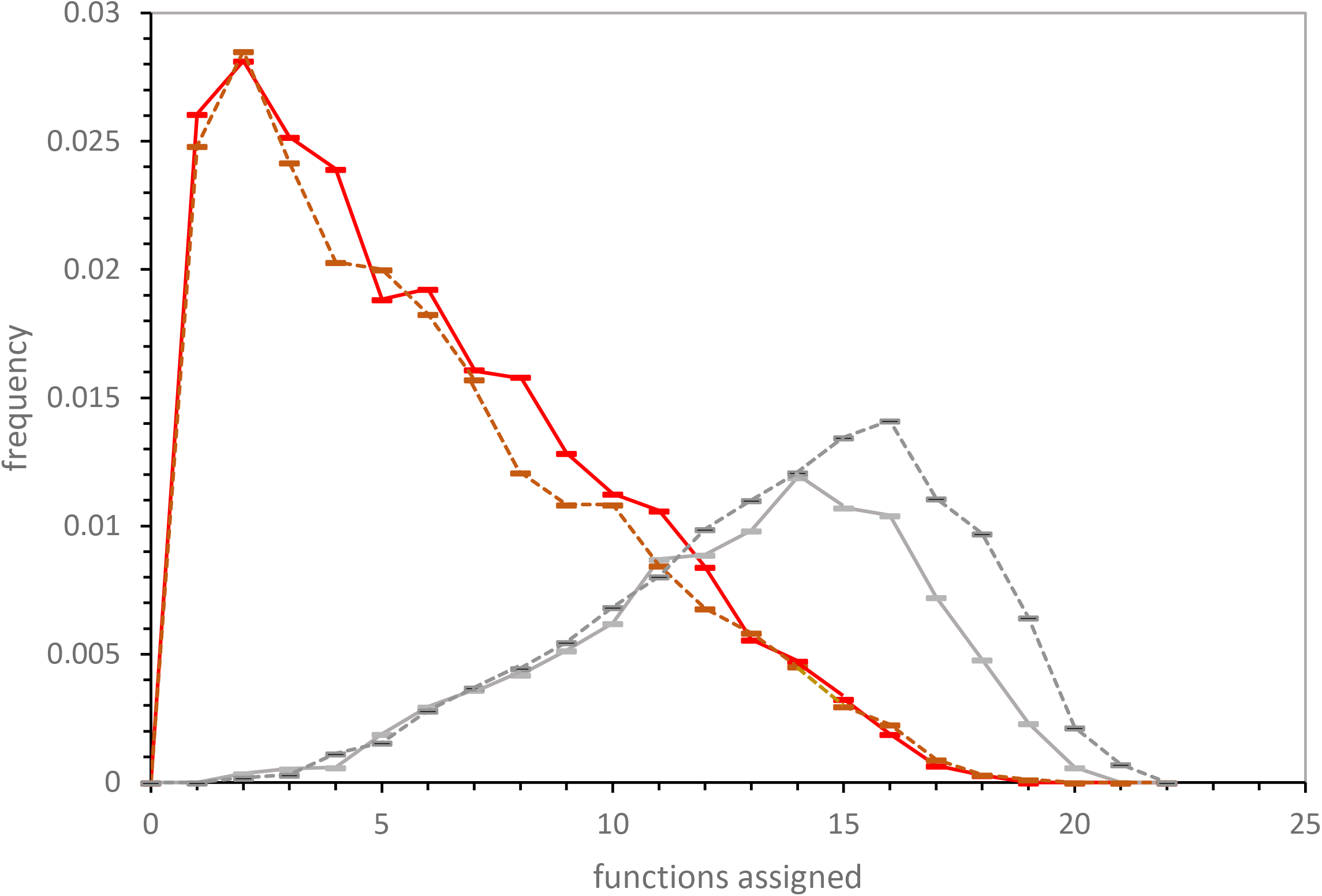
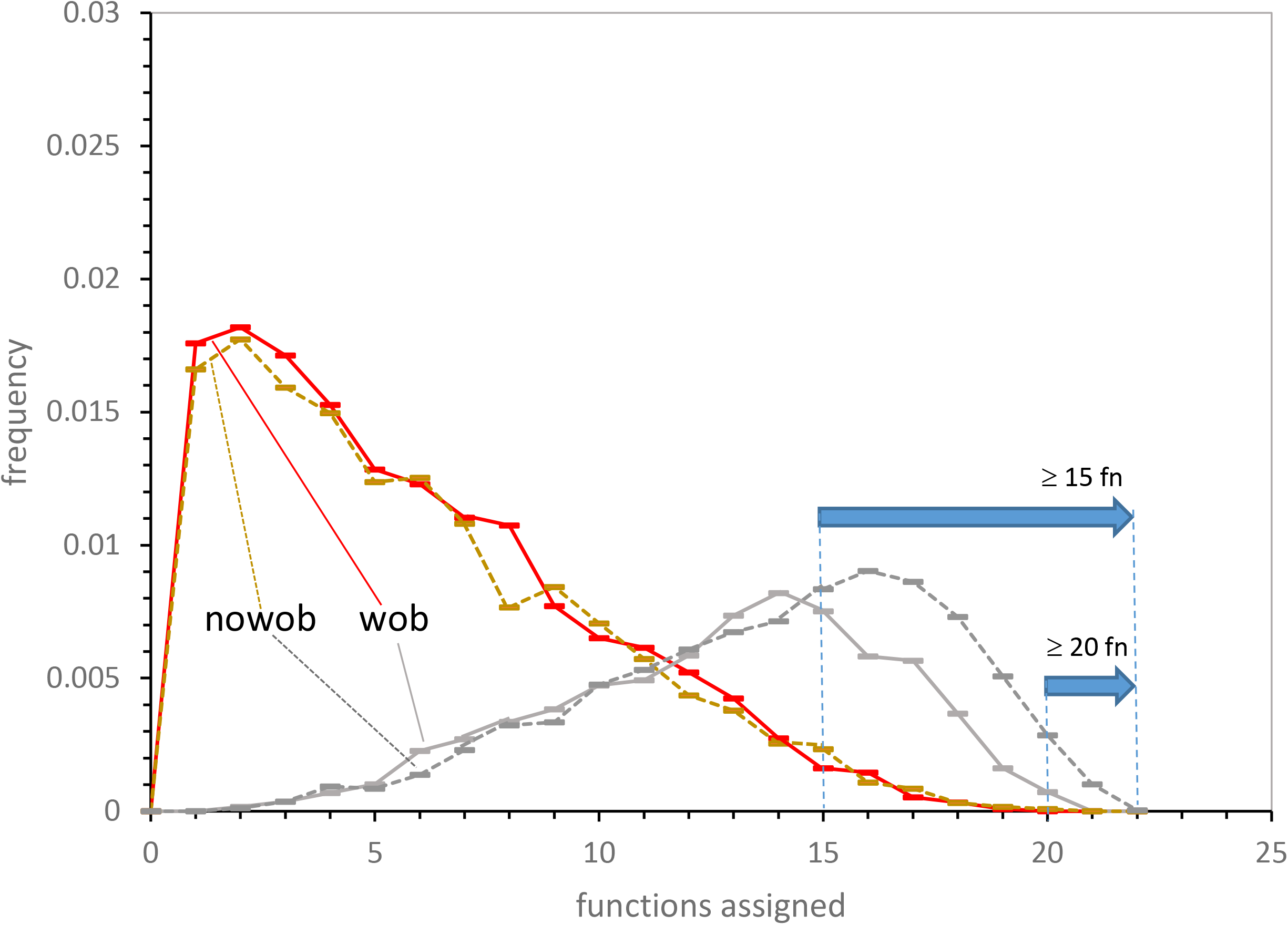
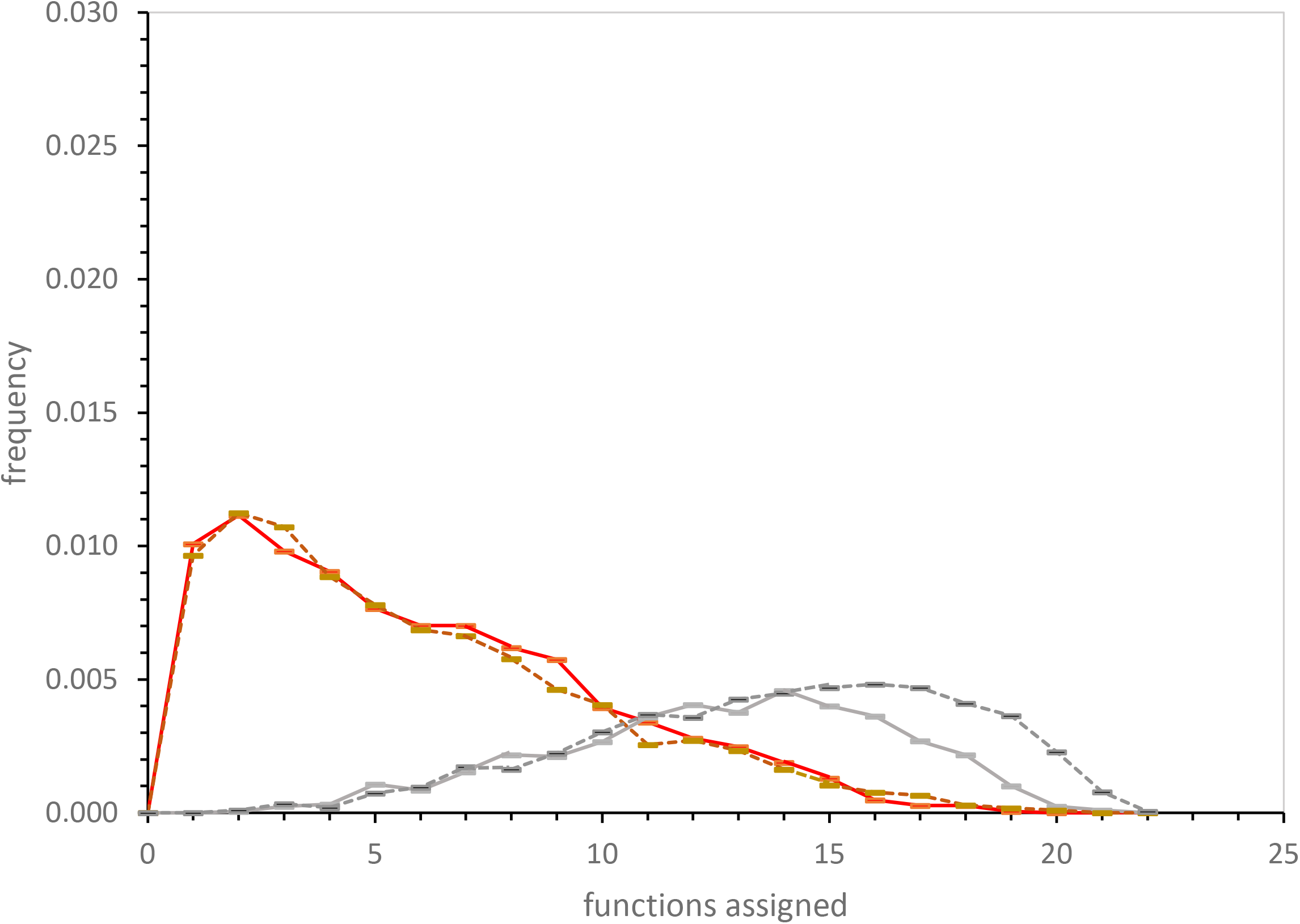
Early wobble specifically disrupts fusion to more complete, SGC-like codes. **Figure 4A** – Fused and unfused active codes are distinct. Mean environmental frequencies versus number of encoded functions in 1000 environments after 300 passages. Pwob = 0.0, and other probabilities/passage as in Fig. 1. Unfused codes (unfus) are red, successful fusions (fused) are gray, and total active codes (sum of unfused and active fusions) are blue. **Figure 4B** – Frequencies of fused and unfused active codes versus number of assignments, with (solid lines, Pwob = 0.02) and without (dashed lines, Pwob = 0.0) wobble. From 1000 environments after 200 passages; other probabilities as in Fig. 1. **Figure 4C** – Frequencies of fused and unfused active codes versus number of assignments, with (marked wob, solid lines, Pwob = 0.02) and without (marked nowob, dashed lines, Pwob = 0.0) wobble. From 1000 environments after 300 passages; other probabilities as in Fig. 1. Horizontal arrows mark codes altered by wobble (>= 15 encoded functions) and closest to the SGC (>= 20 encoded functions). **Figure 4D** – Frequencies of fused and unfused active codes versus number of assignments, with (solid lines, Pwob = 0.02) and without (dashed lines, Pwob = 0.0) wobble. From 1000 environments after 500 passages; other probabilities as in Fig. 1.

Total evolving codes can be divided into two groups with differing fusion histories. A mostly younger group has never fused (red, Fig. 4A). Codes with few functions are mostly unfused, and they contribute little or nothing to the most complete codes. The second group has undergone successful fusions (gray, Fig. 4A) and so has gained encoded functions. These fused codes account for almost all near-complete coding (compare blue and gray, Fig. 4A; (Yarus 2022a)). These fused codes are thus the plausible precursors for an SGC. So: the near-complete code peak formed in the absence of wobble (Fig. 3) is populated principally by fused codes from the region where blue (complete) and gray (fused) codes converge (Fig. 4A). We therefore can sharpen Fig. 3’s conclusion: what is wobble’s effect on critical near-complete fused codes?

### A selective wobble effect on near-complete coding

These data are in Fig. 4B (200 passages), 4C (300 passages) and 4D (500 passages). These codes become competent early in the fusion era (200 passages), pass through the peak of completeness (300 passages) to environments likely to be past the point at which an SGC-candidate would have been selected (500 passages). Fig. 4’s distributions each show a distribution of coding completeness, paralleling the frequency vs assignments presentation of Fig. 4A. However, Fig. 4B, 4C and 4D present only data for active unfused and fused codes, doubled so non-wobbling (dashed lines) and early wobbling populations (solid lines) can be compared.

The similarities in these figures strengthen conclusions below about the likely path of code evolution. Fig. 4B, 4C and 4D all have the same ordinate, so that the decrease in fraction of total codes as unsuccessful code fusions accumulate is evident in Fig. 4’s successive panels. Code fates across the peak of complete codes can be described with one statement: there is little reproducible change among unfused codes if early wobble occurs. The unfused, tending to few assignments, are mostly unchanged by wobble.

Fused codes differ from unfused. In fact, fusion is nearly universal for codes with >= 20 functions at 200, 300 and 500 passages (Fig. 4). Inhibition by early wobble is seen throughout the coding completeness peak (Fig. 3). Early wobble selectively quenches production of codes with >= 15 functions. But this inhibition becomes more extreme if more complete encoding is required. At 200 passages, successful fusions with >=20 functions are decreased 4.4-fold by early wobble (Pwob = 0.02, Fig. 1), at 300 passages decreased 5.5-fold and at 500 passages decreased 8.8-fold (Fig. 4B, 4C and 4D). Selection of a complete SGC from almost-complete codes will be slowed by wobble, and SGC appearance will be relatively slower the later selection occurs (Fig. 2, 3, 4).

### Wobble and accurate assignment

Complete assignment of SGC-like functions has so far been emphasized. But there is a parallel wobble effect on code accuracy. Fusing codes with differing assignments for the same triplet produces ambiguous translation, making the ambiguous fusion less fit. Selection presumably eliminates these discordant codes (Yarus 2022a). Successful fused codes are therefore more homogeneous than their unfused parents, converging on coding assignments common in the initial unfused codes. Convergence to a unique SGC is plausible.

In Fig. 5, wobble coding accuracy is explored in 1000 environments at 300 passages, representing the most likely time for a near-complete code (Fig. 3). Only active, unfused coding tables are represented. In order to summarize the wobble effect, the entire region altered by early wobble is averaged (Fig. 4, >= 15 functions encoded). Such averages usefully summarize distributions; however, note that such distributions extend to near-complete codes, some identical to the SGC (Yarus 2022a).

**Figure 5.**
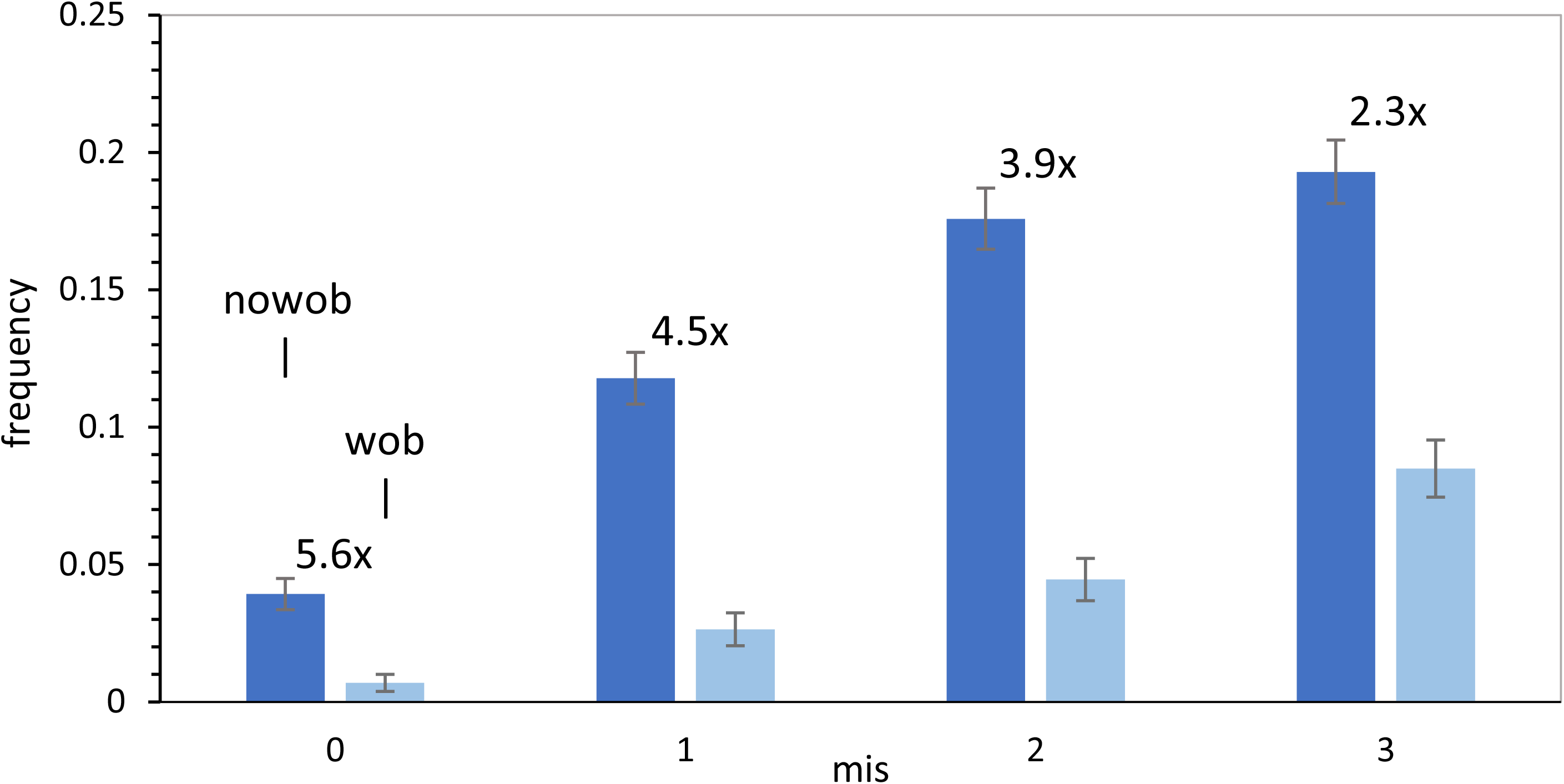
The most complete codes in 1000 environments at 300 passages with (wob) and without wobble (nowob), have different accuracies. Frequency among active codes of the most accurate coding (0, 1, 2 or 3 misassignments - mis0, 1, 2 or 3, respectively), with (Pwob = 0.02) and without (Pwob = 0.0) early wobble. Numbers (e.g., 5.6x) are frequencies for non-wobbling codes divided by frequencies of codes with wobble. Error bars are standard errors for plotted frequencies. Environmental events, save for Pwob, have the probabilities in Fig. 1.

### Early wobble decreases accuracy

Codes with similar assignments are the most likely precursors for selection of the SGC. Fig. 5 concentrates on these; codes with one (mis1), two (mis2) or three (mis3) misassignments, or identical (mis0) to the SGC. Addition of wobble, here and elsewhere, always decreases resemblance to the SGC. That is, wobbling modes (wob) always are further from the SGC than nowob. Among these SGC-like codes, the numbers in Fig. 5 (e.g., 5.6x) are the fold decrease in accurate coding: that is, codes identical to the SGC are decreased 5.6-fold by early wobble. This factor declines as distance from the SGC increases – wobbling codes with three differences from the SGC are 2.3-fold fewer than those with postponed wobble. Wobble is more detrimental, the more accurate an SGC precursor must be. SGC resemblance among fusing coding tables will most readily evolve by avoiding wobble while complete coding is being attained.

## Discussion

### Evolving toward the SGC: completeness

Simplified Crick Wobble (Yarus 2021a) has strong effects if it is probable and becomes frequent early (Fig. 1) during codon assignments in primordial codes. Early wobble can prolong near-complete codon assignment by a factor of 5 or more (Fig. 2). This negative effect is quantitatively explained (Fig. 3) by early wobble’s depressed formation of highly complete codes among those that come from early code fusions. This wobble effect can be further resolved by the distribution of assignments among nascent codes (Fig. 4). Active codes are either yet unfused (red, Fig. 4A) or those successfully fused (gray, Fig. 4A). Many other codes are fused and ambiguous: lost, and not discussed here (Yarus 2022a). Unfused codes dominate coding with few assignments, and fused and summed assignments almost exclusively account for most complete codes (Fig. 4A). When the peak of complete coding (Fig. 3) is examined, wobble changes specifically alter codes completed by fusion (Fig. 4B, 4C and 4D). Consequently, wobble’s obstruction occurs during code fusions. The wobble effect rather specifically depresses formation of more complete codes (≥ 15 functions, marked in Fig. 4C), the more likely precursors of the SGC. Thus early wobble prevents progress toward code completion, probably because assignments extended by wobble present a larger target for conflict between fusing codes. This specifically removes more complete wobbling codes from the fused population, specifically depleting codes with many assigned functions (Fig. 4B, 4C and 4D).

### Evolving toward the SGC: accuracy

Discussion so far is about completing codon assignment, to encode all final functions. But there is a second wobble effect on the accuracy of coding. Code fusion not only sums assignments to yield more complete codes, it also causes codes to converge on assignments shared among fusing codes (Yarus 2022a). In fact, fusion creates a long crescendo of improving adherence to shared assignments. Early wobble obstructs this accuracy-enhancing convergence.

Fig. 5 illustrates this result, showing that among active codes, the four most SGC-like code inventories (mis0, 1, 2 and 3) are depressed by early wobble. Fig. 5 also shows that wobble’s accuracy penalty increases as the SGC is approached. Evolution would likely select a code which was near-complete (Fig. 3) if it could do so without compromising approach to a single code (Vetsigian et al. 2006). Such most homogeneous codes evolve when wobble arises late.

Late Crick wobble was initially suggested because it helps fill the coding table (Yarus 2021a) and join the sections of a fused code (Yarus 2021b). Present work adds that late wobble helps complete SGC coding (Fig. 2, 3 and 4) and also helps converge to a common set of assignments, as in the SGC (Fig. 5).

In a simpler coding system (Yarus 2021c), there was a similarly-timed early optimum for SGC selection, resulting from increase in completeness with time, alongside decrease in accuracy with time. Here, codes evolving via fusion again display an optimal time for SGC selection, but with a newfound optimum due to completeness (Fig. 3, 4C) alone.

### Accurate modern wobble requires multimolecular RNA mechanics

To prevent wobble at first and second codon positions, elaborate multimolecular structures are required. For example, most mutations of the nt 27-43 base pair at the top of the tRNA^Trp^ anticodon hairpin increase first-position wobble, with effects up to 40-fold (Schultz and Yarus 1994). Mutations throughout central structures of tRNA^Ala^ increase both first and second codon position wobble (Shepotinovskaya and Uhlenbeck 2013), as well as improper third position wobble (Olejniczak and Uhlenbeck 2006). Thus, a specific tRNA structure has likely evolved to limit first, second and third position wobble. As emphasized in previous exposition, there are also multiple tertiary ribosomal RNA structure checks on the shape of the codon-anticodon complex (Ogle et al. 2001; Rozov et al. 2016). Thus, to complement full coding, complete coding, and accurate coding arguments made here, there is a compelling structural argument that accurate third position wobble would have been adopted late, after a sophisticated translation apparatus had evolved to edit simple base-pairing. Using the structural argument, earlier calculations have implemented wobble as a late event (Yarus 2021a, 2022a).

### Origins of code order

Coding table structure here is attributable to a small number of captures of related codons (via Pmut: (Yarus 2021a)) and to convergence in assignments due to fusion (Yarus 2022a), based on extension from a subset (Ardell and Sella 2002; Massey 2019) of ancient stereochemical assignments due to amino acid:RNA interactions (Yarus et al. 2005; Yarus 2017). Such stereochemical relations may persist today in regulatory roles, for example, in modern RNA-binding proteins (Kapral et al. 2022). However, this should not be read as an argument against selection against errors (Haig and Hurst 1991; Freeland and Hurst 1998) or against a code co-evolving with amino acid synthesis pathways (Higgs and Pudritz 2009; Wong 1975; Taylor and Coates 1989; Di Giulio 2008). Instead, the contrary seems more plausible: a code fused from diverse origins is consistent with varied sources of order in assembled fragments. Diverse SGC ordering principles are also consistent with varied physicochemical patterns, e.g., within code columns ((Yarus 2021b, 2021c).

### Selection of the historical code

Fusion of nascent partial codes notably offers a persistent series of SGC-like codes, termed the ‘crescendo’ (Yarus 2022a), Fig. 4, 5), for selection as the historical Standard Genetic Code. But what bridged the gap between these long-lasting, varied, SGC-like, late-wobbling precursors and the historical SGC? Because of the crescendo, the final selection plausibly was undemanding – it was a ‘least selection’ (Yarus 2022b) from a highly fit subset within a broad code distribution (Yarus 2021a, Fig. 4). About 4% of active non-wobbling codes, >= 15 functions, are identical to the SGC at 300 passages (Fig. 5): this abundance seems favorable for subsequent SGC emergence.

Selections for favored characteristics are of different kinds (reviewed in Yarus 2022b). The more effective is truncation selection: accepting only entities better than a threshold (Crow and Kimura 1979). Truncation, uniquely, makes improvements of arbitrary size, given large populations. Further, there is a subtler form of truncation that is frequent, sometimes called an ‘evolutionary radiation’ (Yarus 2022b). In such a radiation, minority possessors of some quality become a dominant biota. All radiations are truncations also, thereby associated with rapid evolution. It seems plausible that more complete codes described here supported faster growth based on more varied, accurate protein synthesis. Accordingly, truncation selection in a translation-sustained radiation is a plausible founding event for the SGC.

### Biology as anthology

Other similar paths to biological complexity appear likely. Functional parts arising separately, selectively combined after fusion, genetics or horizontal gene transfer, may provide a general route to multipart, multifunctional competence. This kind of multiply-selected evolutionary progress is deeply characteristic of biological systems. Such grand unions perhaps define biology, including evolution by natural selection as an example. Accordingly, living systems are those that preserve and readily conjoin advances arising separately (compare McKay 2004).

## Methods

### Calculation

Evolutionary time is calculated in passages. In one passage: the model (Yarus 2021a) allows each existing coding table to do one and only one of the following, with the probability Pnnn: make a new assignment (Pinit) of a random function (Prand) or an SGC function and if wobble has begun (Pwob) also assign the additional simple wobble (only third position G:U and U:G pairs allowed; Yarus 2021a) triplet, or an assignment may decay (Pdecay), or an occupied triplet may capture an unassigned neighbor related by a single mutation (Pmut) and assign its existing function or alternatively, an amino acid with a closely related polar requirement (Woese et al. 1966, Mathew and Luthey-Schulten 2008). At each passage, there is a probability that persistent wobble coding will arise (Pwob). Because all events have finite probabilities, a passage can also see no change at all (Yarus 2021a).

Coding tables arise and passage in an environment (Yarus 2022a): at each passage, a new coding table can appear (Ptab). After multiple coding tables exist in an environment (possibly tens or hundreds ultimately exist), two randomly chosen tables can fuse their coding assignments (Pfus). If the fused codes have (a) conflicting codon assignment(s), both fusing codes do not evolve further. These unsuccessful fusions are usually ignored by including only active tables (unfused plus successfully fused) in summary calculations. Fusion initially increases more-than-linearly with time (Yarus 2022a) because of the accumulation of possible fusion partners (via Ptab). However, later coding evolution (e.g, Fig. 3) occurs in a pseudo-steady state with a nearly invariant number of still-evolving coding tables.

A sufficient number of environments are followed (possibly hundreds or thousands) to get a precise idea of the effect of changes (that is, to get a sufficiently small standard error for some mean environmental quantity). Evolution can stop at varied points: for example, when one coding table in the environment becomes complete or near-complete. Full details of coding history (including coding tables) in each environment can be recorded, and potentially available for analysis. But usually, only a quantity from the most complete codes, or the average of all environments at a given passage, are of interest. Calculated output is therefore limited and selected to emphasize data applicable to a current question. Examples of code in Pascal (executed in the developmental environment; Lazarus v 2.2.ORC1, Free Pascal 3.2.2 under 64-bit Windows 10) and of analysis in Microsoft Excel (2016) are available on request.

### Assignments

Pinit is reduced here to ¼ that initially used for calculations (Yarus 2021a), so evolutionary time in passages is slowed, very approximately, four-fold. However, calculations assume one and only coding event per passage, so a slowed Pinit yields a more accurate estimate of kinetics. However, Pinit here is the same as in initial analyses of fusion (Yarus 2021b, 2022a) and those results should be quantitatively similar.

## Notes

### Competing Interest Statement

The authors have declared no competing interest.

